# Nuclear RNA-acetylation can be erased by the deacetylase SIRT7

**DOI:** 10.1101/2021.04.06.438707

**Authors:** Pavel Kudrin, David Meierhofer, Cathrine Broberg Vågbø, Ulf Andersson Vang Ørom

## Abstract

A large number of RNA modifications are known to affect processing and function of rRNA, tRNA and mRNA ^1^. The N4-acetylcytidine (ac4C) is the only known RNA acetylation event and is known to occur on rRNA, tRNA and mRNA ^2,3^. RNA modification by acetylation affects a number of biological processes, including translation and RNA stability ^2^. For a few RNA methyl modifications, a reversible nature has been demonstrated where specific writer proteins deposit the modification and eraser proteins can remove them by oxidative demethylation ^4–6^. The functionality of RNA modifications is often mediated by interaction with reader proteins that bind dependent on the presence of specific modifications ^1^. The NAT10 acetyltransferase has been firmly identified as the main writer of acetylation of cytidine ribonucleotides, but so far neither readers nor erasers of ac4C have been identified ^2,3^. Here we show, that ac4C is bound by the nucleolar protein NOP58 and deacetylated by SIRT7, for the first time demonstrating reversal by another mechanism than oxidative demethylation. NOP58 and SIRT7 are involved in snoRNA function and pre-ribosomal RNA processing ^7–10^, and using a NAT10 deficient cell line we can show that the reduction in ac4C levels affects both snoRNA sub-nuclear localization and pre-rRNA processing. SIRT7 can deacetylate RNA *in vitro* and endogenous levels of ac4C on snoRNA increase in a SIRT7 deficient cell line, supporting its endogenous function as an RNA deacetylase. In summary, we identify the first eraser and reader proteins of the RNA modification ac4C, respectively, and suggest an involvement of RNA acetylation in snoRNA function and pre-rRNA processing.

## Introduction

The number of known RNA modifications exceeds 150 distinct types ^1^. Implication of RNA modifications in biological processes and disease suggests their importance in regulatory pathways and as candidate drug targets ^11^. One of the best characterized RNA modifications is N6-methyladenosine (m^6^A) that is widely accepted to be reversible in nature. The impact of m^6^A on gene expression and RNA processing is exerted through the modification-mediated interaction with RNA-binding proteins. Writer proteins, proteins depositing the specific RNA modifications, have been identified for m^6^A and other RNA modifications, and in some cases also eraser proteins, removing the RNA modifications in a reversible manner, have been identified ^12^. Most RNA modifications, as e.g. pseudouridine, are considered irreversible ^13^.

The N4-acetylcytidine modification (ac4C) has been shown to be present in tRNA and 18S rRNA ^14^, and recently also demonstrated to be present at mRNA with a proposed role in mRNA stability and translation fidelity ^2^. In eukaryotes, neither readers nor erasers of ac4C have been identified to date, and the only known writer is the nucleolar protein NAT10. Knocking out NAT10 in human cancer cells results in an 80 per cent reduction of global ac4C levels ^2^, and a similar effect is observed in archaea in response to deletion of the NAT10 homolog Tk0754 ^3^, suggesting that additional ac4C writers might exist. Under normal growth conditions NAT10 localizes to the nucleolus, a cellular compartment organized around rDNA genes clustered into nucleolar organizing regions (NOR) ^15^. The main task of nucleoli is rRNA transcription, processing and ribosome subunit assembly. In mammals the primary pre-rRNA transcript (47S pre-rRNA) consists of the precursors to 18S, 5.8S and 28S rRNAs and internal and external transcribed spacers (ITS1-2; 5’ ETS; 3’ ETS), and is processed sequentially ^16^. The nascent 47S pre-rRNA transcript is a target for posttranscriptional processing conducted by snoRNP complexes guided by snoRNAs. Aside from spacer trimming rRNA processing also includes 2’-O-ribose methylation (2’-OMe) and pseudouridylation (Ψ) in a site-specific manner directed by box C/D and box H/ACA snoRNPs respectively ^17^. Interestingly, the yeast homolog of NAT10, Kre33, positions ac4C on 18S rRNA under the guidance of box C/D snoRNAs ^18^. Here, we show that the nucleolar protein NOP58 binds to ac4C and identify the SIRT7 as the first protein that can remove ac4C from RNA, suggesting that RNA acetylation is a reversible process that can affect RNA localization and processing.

## Results

To identify protein binders recognizing the ac4C modification, we *in vitro* synthesized a biotinylated 76 nts RNA. The RNA contains 24 C nucleotides in various sequence contexts and was synthesized in the presence of unlabeled A, U and G ribonucleotides and increasing concentrations of ac4C modified C (0, 50 and 100 per cent ac4C over C, respectively). The *in vitro* synthesized RNAs were incubated with total cellular lysate, or nucleolar lysate, from HeLa cells deficient of NAT10 (Figure 1a). We eluted bound proteins from RNA using excess biotin and RNase, and subjected purified proteins to liquid-phase mass spectrometry (LC/MS). Here, we identify the NOP58 protein enriched by the ac4C labeled RNA compared to the control non-acetylated RNA. We validate the interaction between ac4C and NOP58 with western blotting using a NOP58-specific antibody, confirming an ac4C-dependent binding of NOP58 to RNA (Figure 1b).

**Figure 1.**
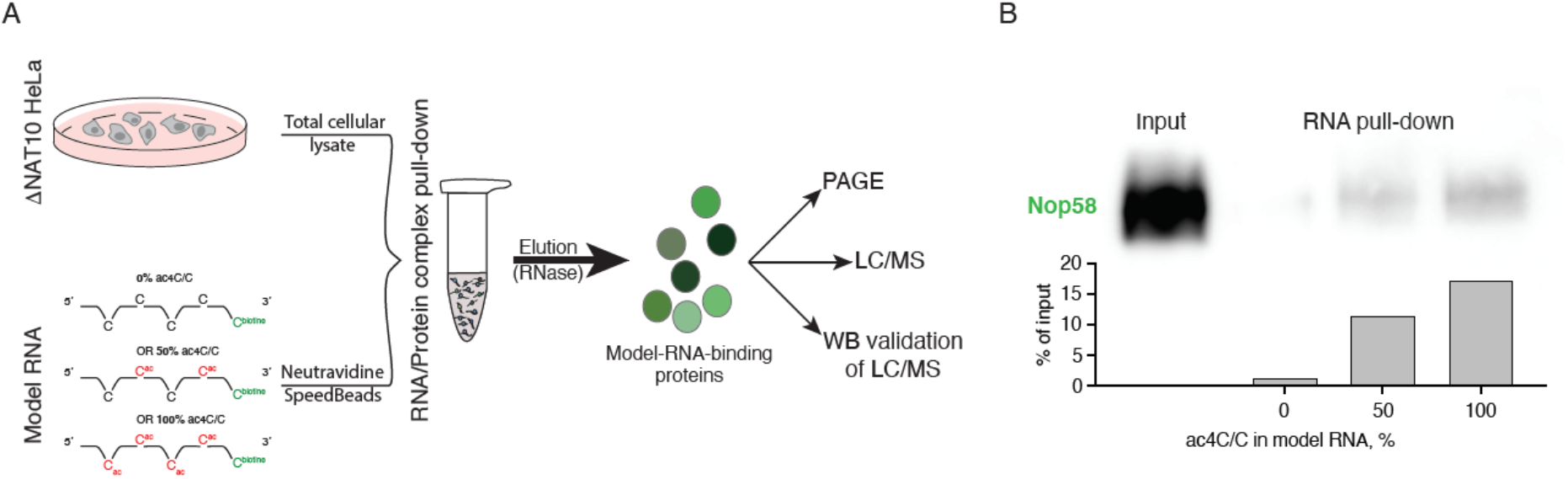
Identification of NOP58 as ac4C-binding protein. a) Overview of the experimental approach; b) Immunobloting of NOP58 from ac4C-enriched RNA-binding-protein pull-down using *in vitro* synthesized RNA with 0, 50 and 100 percent ac4C/C, respectively. Quantification of the protein bands is shown in the lower panel.

NOP58 is a nucleolar protein that binds to snoRNAs and mediates their nuclear to nucleolar shuttling as well as their recruitment to pre-rRNA. Both processing of the 47S rRNA and modification of individual subunits are affected by snoRNAs with involvement of the NOP58 protein ^7,8^. To assess whether ac4C affect snoRNA localization we purified RNA from fractionated nuclei and nucleoli from the NAT10 deficient cell line (Supplementary Figure 1). Here, we observe that snoRNAs tend to accumulate in the nucleolus in the NAT10 deficient cell line, supporting a role for ac4C in snoRNA sub-nuclear localization (Figure 2a). As snoRNAs can affect both pseudouridylation and 2′-O-methylation of 28S and 18S rRNAs ^17^ we assessed RNA modification levels on 28S and 18S rRNA subunits of all known RNA modifications by RNA mass spectrometry to see if a reduction of ac4C affects the levels of other RNA modifications. We find 80 per cent reduced ac4C levels on both 28S (Figure 2b) and 18S (Figure 2c) subunits, but no changes to pseudouridine or 2′-O-methylation levels (Figure 2b-c), or any of the other assessed modifications (Supplementary Figure 2), suggesting that NAT10 deprivation does not affect other rRNA modifications than ac4C. We notice on the bioanalyzer spectra that the pre-RNA 45S/47S band is more intense in NAT10 deficient cells than in WT HeLa, and also that the 18S rRNA band is less intense when ac4C levels are reduced (Figure 2d), suggesting that pre-rRNA processing is less efficient in the absence of ac4C modified RNA. We validate this effect on 45S/47S processing by qPCR using primers spanning one of the processing sites, and we show that both 47S pre-rRNA and 45S pre-rRNA transcripts accumulate in the absence of ac4C (Figure 2e). Data in Figure 2a and 2e are normalized to MALAT1, a nuclear long non-coding RNA where the localization is not affected by the depletion of NAT10 (Supplementary Figure 3).

**Figure 2.**
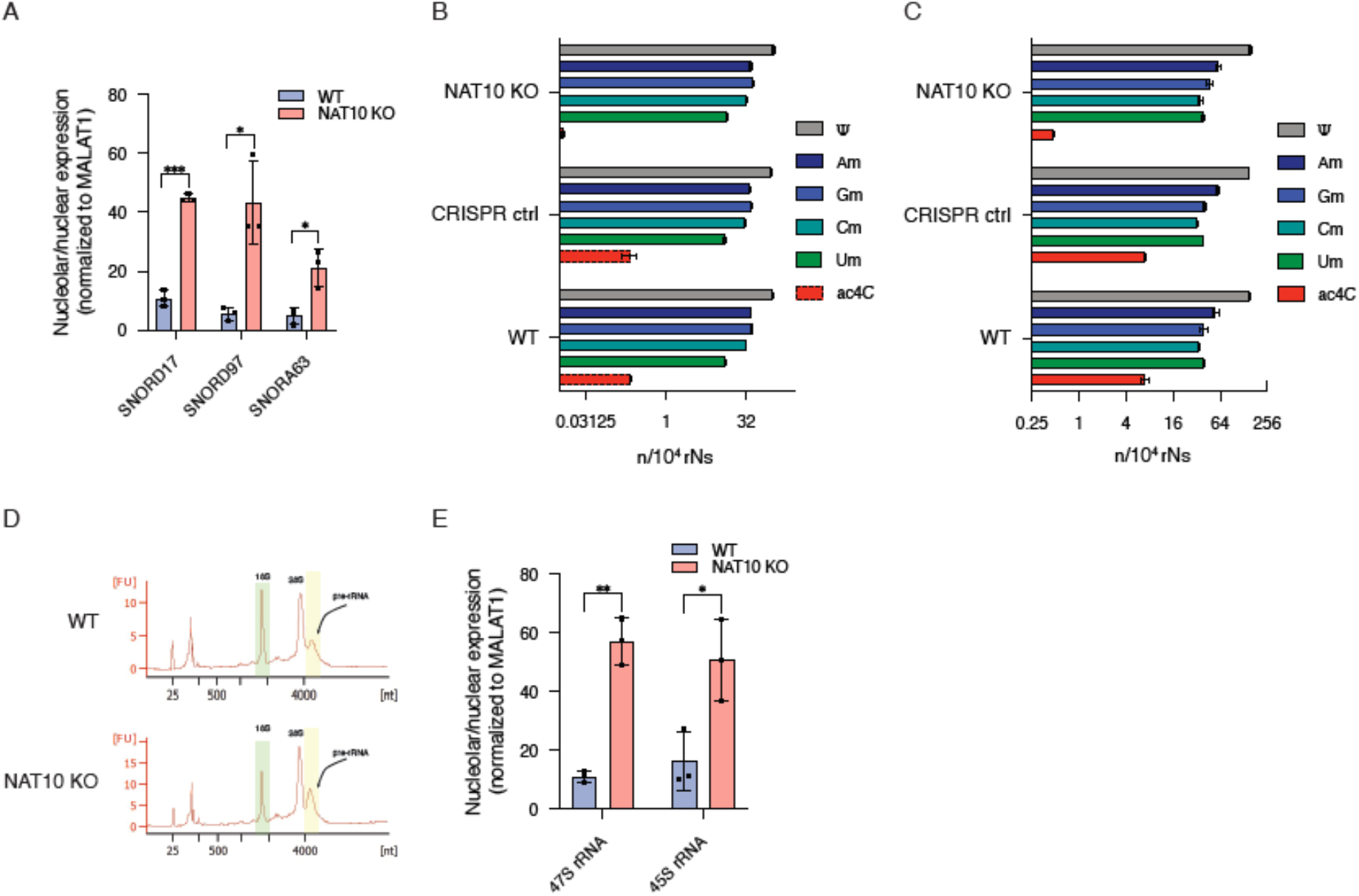
ac4C is important for pre-rRNA processing. a) snoRNA localization in WT and NAT10 KO cells validated by qRT-PCR; b) RNA MS analysis of 28S rRNA for RNA modifications ac4C, Ψ and 2’-OMe. Dashed ac4C profile indicates that ac4C was not previously annotated in 28S rRNA; c) RNA MS analysis of 18S rRNA for RNA modifications ac4C, Ψ and 2’-OMe; d) Representative Bioanalyzer spectrum of the distribution of RNA species in WT and NAT10 KO cells. Regions for 18S rRNA and 47/45S pre-rRNA are highlighted in green and yellow respectively; e) 47S and 45S pre-rRNA nucleolar enrichment in WT and NAT10 KO cells validated by qRT-PCR. n=3 independent experiments for panels a and e and n=2 independent experiments for panels b and c. Data are shown as average ± SD with individual data points shown. Significance is determined using one-tailed Student’s t-test (*p<0.05, **p<0.01, ***p<0.005).

To identify putative deacetylases we looked for protein candidates that can interact with NOP58 in the nucleolus and has an involvement in pre-rRNA processing, as both NAT10 and NOP58 have been shown to affect pre-rRNA processing in a nucleolar screen of pre-rRNA processing factors ^10^. A number of protein deacetylases are known, including histone deacetylases (HDAC) and Sirtuins ^19^. Only one of these proteins has been shown to affect pre-rRNA processing, the SIRT7 protein ^10^. SIRT7 interacts with NOP58 ^20^ and has been shown to be an RNA-activated protein deacetylase ^9^. The RNA binding properties of SIRT7, its known interaction with NOP58, the effects on pre-rRNA processing and its nucleolar localization are all pointing towards SIRT7 as a putative RNA deacetylase. We set up an *in vitro* deacetylation assay using *in vitro* transcribed ac4C-containing RNA and recombinant SIRT7 followed by dot blot and probing with an ac4C-specific antibody (Figure 3a). We show that SIRT7 is able to deacetylate ac4C within RNA *in vitro* and that this deacetylation appears to be NAD^+^ independent (Figure 3b). To determine whether SIRT7 has an endogenous impact on ac4C we quantified ac4C in RNA from a SIRT7 knock-out cell line using RNA mass spectrometry. Here, we observed a slight decrease in the global ac4C levels, whereas a m2,2,7G-modified snoRNA fraction had increased ac4C levels (Figure 3c), suggesting that SIRT7 acts as an RNA deacetylase on a subset of ac4C-modified RNA species inside the cell.

**Figure 3.**
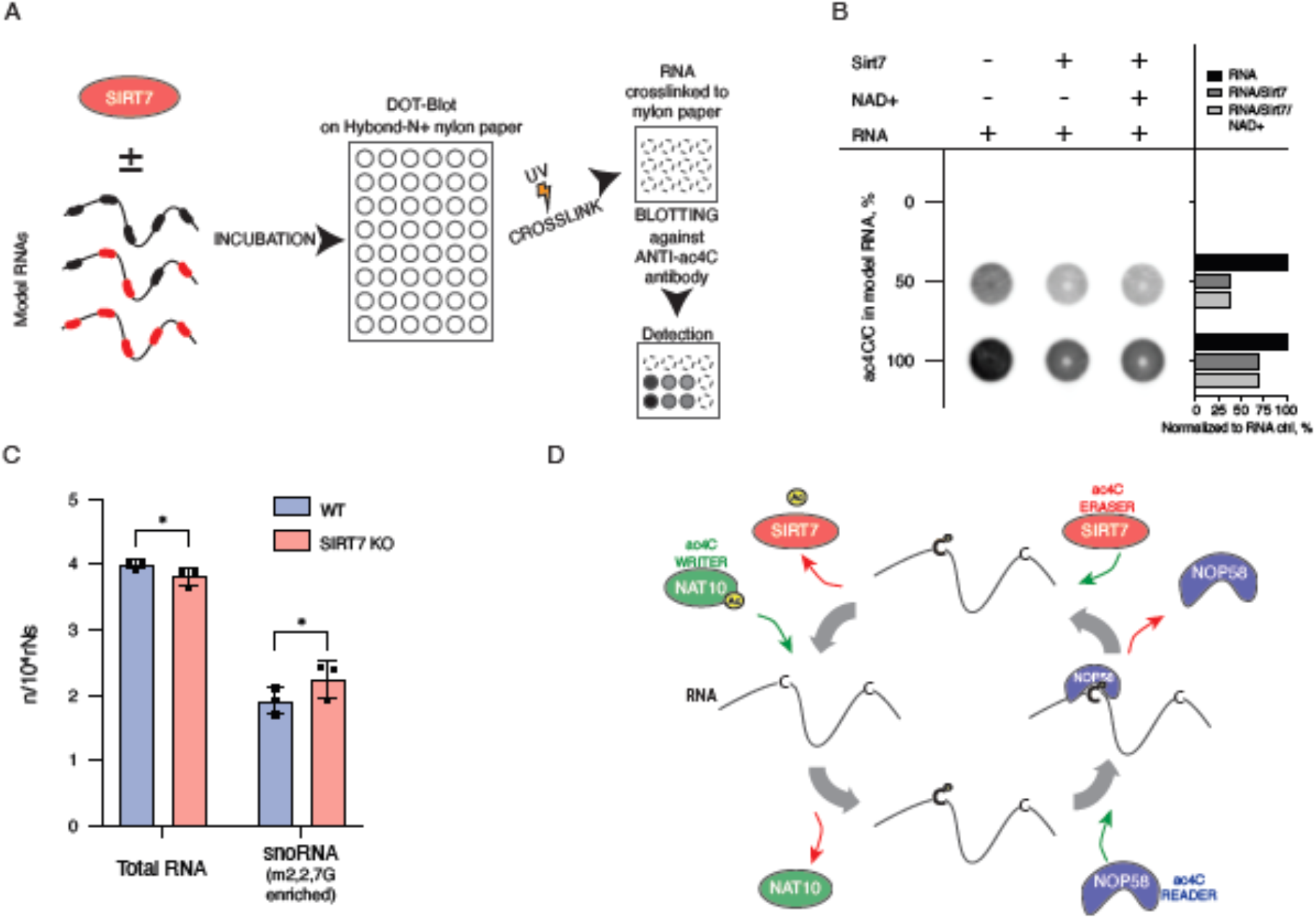
SIRT7 is a nuclear ac4C deacetylase. a) Experimental approach of the *in vitro* deacetylase assay; b) SIRT7 removes ac4C RNA acetylation *in vitro*. Immunoblotting using anti-ac4C antibody to detect ac4C levels of synthetic RNA preincubated with 1 μg recombinant SIRT7 with or without NAD+. Quantification of signal is shown in the right panel; c) ac4C levels in endogenous total RNA and m2,2,7G-enriched snoRNA from WT and SIRT7 knock-out cells determined by RNA mass spectrometry. n=3 independent experiments. Data are shown as average ± SD and individual data point are shown. Significance determined using one-tailed Student’s t-test (*p<0.05); d) Schematic overview of the proposed dynamic acetylation and deacetylation of cytidine with writer NAT10, binder NOP58, and eraser SIRT7.

## Discussion

The identification of the first ac4C reader (NOP58) and ac4C eraser (SIRT7) shows that RNA acetylation is reversible. Deacetylation by SIRT7 is the first demonstration of RNA modification reversal by a different mechanism than oxidative demethylation, and expands the repertoire of RNA modifications that are dynamically regulated in cells. SIRT7 has been shown to bind nucleotides and to be particularly activated by RNAs ^9^, in line with our findings that SIRT7 acts as an RNA deacetylase. Previous work solved difficulties in making SIRT7 work *in vitro* by adding rRNA and tRNA to the assay ^9^ suggesting that RNA is required for SIRT7 function. In the presented data the deacetylation effect of SIRT7 on synthetic RNA is not complete which could be due to the high occurrence of C’s in the RNA sequence (24 C’s in the RNA sequence). As no consensus site for RNA acetylation is known it is possible that we have included artificial sites in the sequence that are not accessible to SIRT7.

In Figure 3d we summarize the reversible and dynamic deposition of ac4C onto RNA by NAT10 and SIRT7. SIRT7 is the only protein deacetylase with a nucleolar function in pre-rRNA processing, yet other proteins could work as RNA deacetylases in distinct cellular compartments where other RNA types are predominant, as *e.g.* mRNA in the cytoplasm. While NAT10 depletion by CRISPR-mediated knock-out clearly removes most of ac4C from all types of RNA there is still 20 per cent left, suggesting that other RNA acetyltransferases could exist as well, and underlines that our understanding of dynamic RNA acetylation is not yet complete. Here, we identify SIRT7 as the first RNA ac4C deacetylase and suggest a functional role for ac4C in snoRNA function and pre-rRNA processing, expanding the molecular insight into dynamic RNA acetylation and suggests a novel function for ac4C in non-coding RNA biogenesis and function.

## Methods

### Tissue culture

If not stated otherwise, HeLa and HEK293F cells were grown in Dulbecco’s Modified Eagle’s medium (DMEM; Gibco, #41966-029 and #41965-039 respectively) supplemented with 10% Fetal Bovine Serum (FBS; Gibco) and 1% Penicillin/Streptomycin (P/S; Gibco) at 37°C with 5% CO_2_ until 90% confluency. Cells were collected by trypsinization with 0.05% Trypsin-EDTA (Gibco). When needed the purification of nuclei was performed as per Conrad and Ørom ^21^ and nucleoli were isolated as per Li and Lam ^22^. Fractionation efficiency assessment by microscopy was performed with Zeiss AxioVert 200M microscope. Consecutive total, nuclear or nucleolar RNA purification was performed using TRIzol (Invitrogen) according to manufacturer’s instructions and RNA preparation quality analyzed with 2100 Bioanalyzer (Agilent).

### In vitro transcription of ac4C modified RNA

T7 promoter containing double stranded DNA was used as a template for in vitro transcription with HighYield T7 mRNA Synthesis Kit (ac4CTP) (Jena Bioscience) (sense strand 5’ GTACGGTAATACGACTCACTATAGGGAGTGGTCTACACACATGACAGAATGGGGCAGGTCCGTAATCGGTTGCAGAGCGGTTACCGATCTCATCGC 3’ and antisense strand 5’ GGCCGCGATGAGATCGGTAACCGCTCTGCAACCGATTACGGACCTGCCCCATTCTGTCATGTGTGTAGACCACTCCCTATAGTGAGTCGTATTACC 3’). CTP and ac4CTP substrates were used in different combinations to obtain the certain level of acetylated cytidines within RNA. Purified RNA was subjected for 3’ end biotinylation with Pierce™ RNA 3’ End Biotinylation Kit (Thermo) resulting in 76 nt long RNA of the following sequence 5’GGCCGCGAUGAGAUCGGUAACCGCUCUGCAACCGAUUACGGACCUGCCCCAUUCUGUCAUGUGUGUAGACCACUC-C(biotine)3’ with either 0%, 50% or 100% of Cs acetylated.

### Purification of ac4C binding proteins

50 μL of Neutravidine SpeedBeads (Sigma) beads per reaction were equilibrated in buffer A (20 mM Tris-HCl pH 7.4, 1M NaCl, 1 mM PMSF, PI cocktail, Superase-in (1 U/ml) (Thermo), 1 mM EDTA) followed by addition of 50 pmol of biotinylated synthetic RNA with defined level of acetylated Cs. After the incubation on rotator at RT for 1 h the beads were washed three times and resuspended in buffer B (20 mM Tris (pH 7.4), 50 mM NaCl, 2 mM MgCl2, 0.1% Tween™-20). During the incubation step the HeLa total cell lysate was prepared by lysing freshly collected HeLa cells in RIPA buffer (Sigma) (1 ml per 15 cm plate) supplemented with 1: 100 protease inhibitor cocktail (Sigma) and 1 mM PMSF on ice for 15 min followed by sonication and another 15 min on ice. Cell debris was removed by centrifugation at 4 °C for 10 min at ≥10000 g. Protein concentration was determined by Bradford assay. 100 μg of HeLa lysate per reaction was mixed with 1x buffer B, 15% glycerol and RNase-free H_2_O with consecutive addition on RNA-beads mix and incubation at 4°C for 60 min with rotation. The beads were washed three times with Wash buffer (20 mM Tris (pH 7.4), 10 mM NaCl, 0.1% Tween™-20, 1 mM PMSF, 1:100 PI cocktail) and RNA-bound proteins eluted in 28 μL of elution buffer (1 mM biotin in Wash Buffer and 2 μL RNase) by incubation shaking at 37°C for 30 min. Eluted proteins were subsequently analyzed by PAGE, Western Blot and MS.

### Proteomics Sample Preparation and LC-MS/MS Instrument Settings

Samples were delivered in 1x PBS, 0.01% SDS, the pH was adjusted to 8.5 by adding a final concentration of 100 mM Tris, followed by denaturing at 95°C for 10 minutes at 1000 rpm. 4 μg protein of each sample was further processed. Reduction of cysteines was carried out by adding 1.1 μl of 0.1 M tris(2-carboxyethyl)phosphine at 37°C for 30 minutes at 800rpm, alkylation of cysteines similarly by adding 2.5 μl of 0.2 M 2-chloroacetamide. Samples were digested by trypsin (enzyme-protein ratio 1:40) at 37°C overnight, desalted and reconstituted in 2% formic acid and 5% acetonitrile in water prior to injection to nano-LC-MS. For each sample, 1 and 3 μg protein were injected. LC-MS/MS was carried out by nanoflow reverse phase liquid chromatography (Dionex Ultimate 3000, Thermo Scientific, Waltham, MA) coupled online to a Q-Exactive HF Orbitrap mass spectrometer (Thermo Scientific, Waltham, MA), as reported previously ^23^. Raw MS data were processed with MaxQuant software v1.6.10.43 ^24^, runs from the same samples were combined and searched against the human UniProtKB with 75,074 entries, released in 05/2020.

### Validation of ac4C binding proteins and western blot

Putative ac4C binding proteins were subjected to SDS PAGE on Novex Tris-Glycine 4-20% (Invitrogen) gel followed by either silver staining with Pierce Silver Stain Kit (Thermo) or Western blotting against anti-Nop58 (#ab236724, Abcam) and anti-GAPDH (#5174s, Cell Signaling Tech) antibodies. Western blot was developed using Pierce ECL Western Blotting Substrate (Thermo) and imaged with Amersham Imager 600 (GE Healthcare).

### Size-exclusion chromatography of total RNA

Total RNA was fractionated into tRNA, 18S rRNA and 28S rRNA using two dimensions of size-exclusion chromatography (SEC) carried out on an Agilent HP1200 HPLC system with UV detector and fraction collector. The 1^st^ SEC dimension was performed using a Bio SEC-5 1000 Å, 5 μm, 7.8 × 300 mm column (Agilent Technologies, Foster City, CA) and isocratic elution with 100 mM ammonium acetate (pH 7.0) at 500 μl/min for 40 min at 60°C, collecting three fractions containing 28S rRNA, 18S rRNA, and RNAs below 200 nt (‘small RNAs’), respectively. The fractions were lyophilized and the small RNA fraction was reconstituted in 20 μl of water and subjected to a 2^nd^ dimension of SEC using an AdvanceBio SEC 120 Å, 1.9 μm, 4.6 × 300 mm column (Agilent Technologies, Foster City, CA) and isocratic elution with 100 mM ammonium acetate (pH 7.0) run at 150 μl/min for 40 min at 40°C.

### Analysis of isolated RNA species using LC-MS/MS

RNA was hydrolyzed to ribonucleosides by 20 U benzonase (Santa Cruz Biotech) and 0.2 U nuclease P1 (Sigma-Aldrich, Saint-Louis, MO) in 10 mM ammonium acetate pH 6.0 and 1 mM magnesium chloride at 40 °C for 1 hour, then added ammonium bicarbonate to 50 mM, 0.002 U phosphodiesterase I and 0.1 U alkaline phosphatase (Sigma-Aldrich, Saint-Louis, MO) and incubated further at 37 °C for 1 hour. The hydrolysates were added 3 volumes of acetonitrile and centrifuged (16,000 g, 30 min, 4 °C). The supernatants were lyophilized and dissolved in 50 μl water for LC-MS/MS analysis of modified and unmodified ribonucleosides. Chromatographic separation was performed using an Agilent 1290 Infinity II UHPLC system with an ZORBAX RRHD Eclipse Plus C18 150 × 2.1 mm ID (1.8 μm) column protected with an ZORBAX RRHD Eclipse Plus C18 5 × 2.1 mm ID (1.8 μm) guard column (Agilent Technologies, Foster City, CA). The mobile phase consisted of water and methanol (both added 0.1% formic acid) run at 0.23 ml/min, for modifications starting with 5% methanol for 0.5 min followed by a 2.5 min gradient of 5-15 % methanol, a 3 min gradient of 15-95% methanol and 4 min re-equilibration with 5% methanol. A portion of each sample was diluted for the analysis of unmodified ribonucleosides which was chromatographed isocratically with 20% methanol. Mass spectrometric detection was performed using an Agilent 6495 Triple Quadrupole system with electrospray ionization, monitoring the mass transitions 268.1-136.1 (A), 284.1-152.1 (G), 244.1-112.1 (C), 245.1-113.1 (U), 282.1-136.1 (Am), 326.1-194.1 (m^2,2,7^G), 286.1-154.1 (ac^4^C), 258.1-112-1 (Cm), and 259.1-113.1 (Um) in positive ionization mode, and 243.1-153.1 (Ψ) in negative ionization mode.

### qPCR and RNA purification

To validate 45S and 47S rRNA transcripts from previously purified nuclear and nucleolar RNA by qPCR cDNA was synthesized with RevertAid First Strand cDNA Synthesis Kit (Thermo) according to manufacturer’s protocol. Platinum™ SYBR™ Green qPCR SuperMix-UDG (Thermo) and the following primers were used for qPCR:

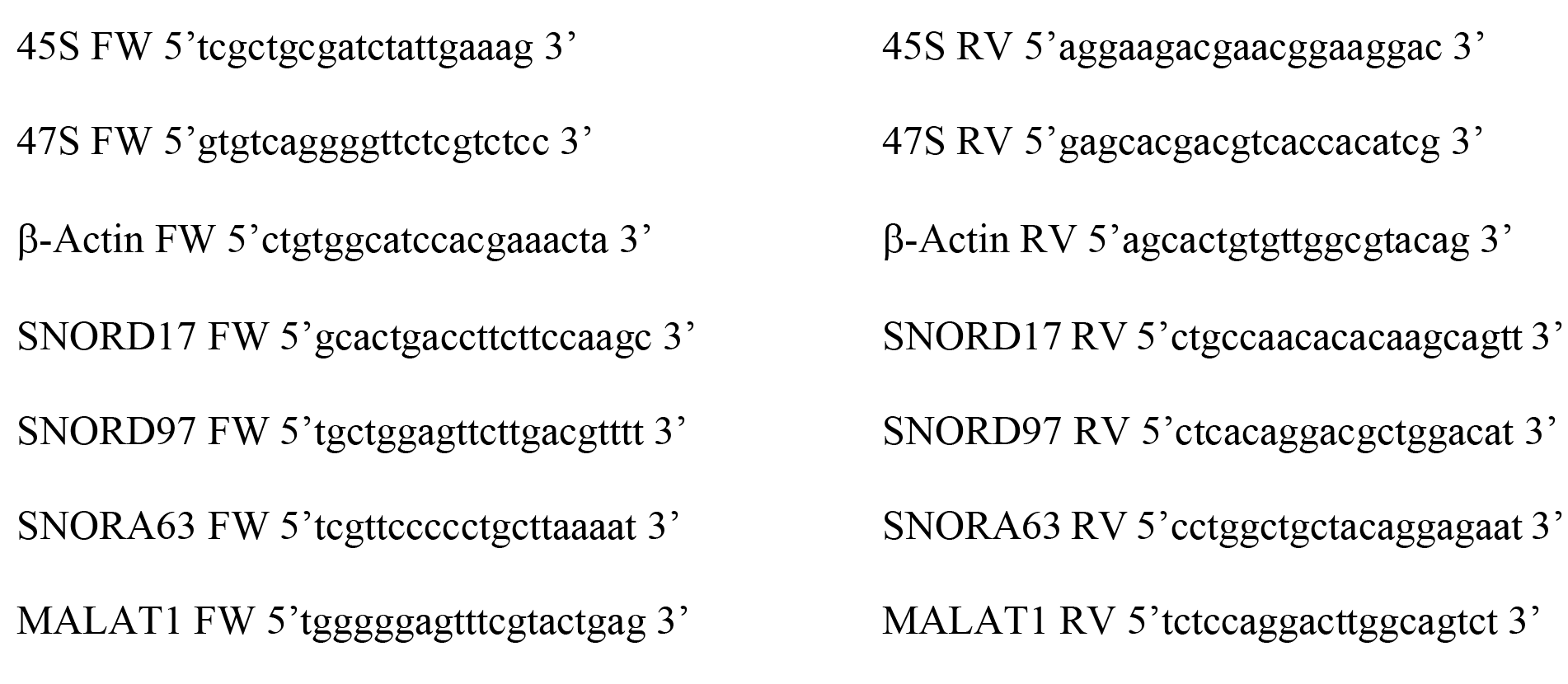

### SIRT7 in vitro deacetylase assay

1 μg Sirt7 protein (#ab104032, Abcam) was preincubated with 50 pmol synthetic RNA containing defined levels of ac4C (0, 50 per cent or 100 per cent ac4C/C) in the following buffer: 10 mM Tris pH 8.0, 2 mM MgCl_2_, 0.2 mM DTT and 10% glycerol either in the presence or absence of 2.5 mM NAD^+^. The reaction was carried out at 30°C for 1h shaking (750 rpm). Using dot-blot manifold the reaction mixture was transferred onto Amersham Hybond-N+ (GE Healthcare) nylon paper and crosslinked at 120 000 μJ/cm^2^. After blotting against anti-ac4C antibody (#ab252215, Abcam) the development was performed using Pierce ECL Western Blotting Substrate (Thermo) and result analyzed with Amersham Imager 600 (GE Healthcare).

### snoRNA purification with m2,2,7G IP

40 μl of Protein G Dynabeads (Invitrogen) in IP buffer (150 mM NaCl, 10 mM Tris-HCl, pH 7.5, 0.1% IGEPAL CA-630 in nuclease free H_2_O) were tumbled with 10 μg anti-m2,2,7G antibody (#MABE302, Sigma) at 4°C for at least 6 hrs. Upon previously purified total RNA addition, RNA-antibody-beads mixture in IP buffer was incubated ON at 4°C with gentle rotation in a final volume of 0.8 mL in protein low-binding tubes. For elution, the beads were resuspended in 1x Proteinase K buffer (100 mM Tris-HCl pH 7.5, 150 mM NaCl, 12.5 mM EDTA, 2% SDS and 120 μg/ml Proteinase K (Invitrogen)) and incubated 1 hr with continuous shaking (1200 rpm) at 37°C. Magnetic separation rack was applied to collect the supernatant. 700 μl of RLT buffer and 1400 μl of 100% ethanol were added to 200 μl of eluted supernatants collected and mixed thoroughly. The mixture was transferred to an RNeasy MiniElute spin column (QIAGEN) and centrifuged at >12000 rpm at 4°C for 1 min. This step was repeated until all sample was loaded to the column. The spin column membrane was washed with 500 μl of RPE buffer once, then 500 μl of 80% ethanol once and centrifuged at full speed for 5 min at 4°C remove the residual ethanol. The m2,2,7G enriched RNA was eluted with 14 μl ultrapure H_2_O. RNA concentration was measured using the Qubit RNA HS Assay Kit as per the manufacturer’s instructions

### Statistics

All statistics are done using Student’s T-test.

## Acknowledgements

We thank Beata Lukaszewska-McGreal for proteome sample preparation and Anna Kusnierczyk for RNA mass spec analysis. We thank Shalini Oberdoerfer for kindly providing the NAT10 KO and the CRISPR control HeLa cell lines and Alejandro Vaquero for kindly providing the SIRT7 KO and parental HEK293F cell line. Work in the author’s lab is funded by the Novo Nordisk Foundation, The Lundbeck Foundation, Danish Cancer Society, Independent Research Fund Denmark, The Carlsberg Foundation to UAVØ and the Max Planck Society to DM.

## Author contribution

PK conceived the experiments, performed the experiments, analyzed data, interpreted data, wrote the manuscript. DM performed mass spectrometry identification of proteins and analyzed data. CBV conceived and performed RNA mass spectrometry experiments and analyzed data. UAVØ conceived the experiments, interpreted data, supervised research, secured funding, wrote the manuscript. All authors read and approved the final version of the manuscript.

