## Supplementary material for "Nuclear RNA-acetylation can be erased by the deacetylase SIRT7": Supplemetary Figures

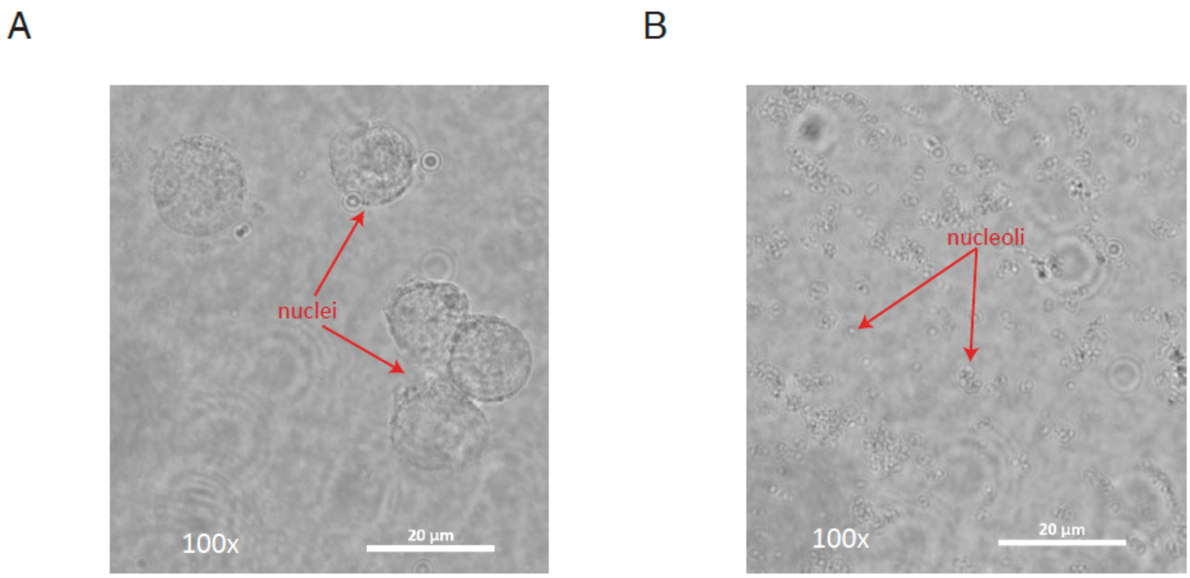


**Figure S1. Bright-field representative images showing cellular fractionation efficiency.** a) Nuclei; b) Nucleoli. 100x magnification, scale bar 20 µm.

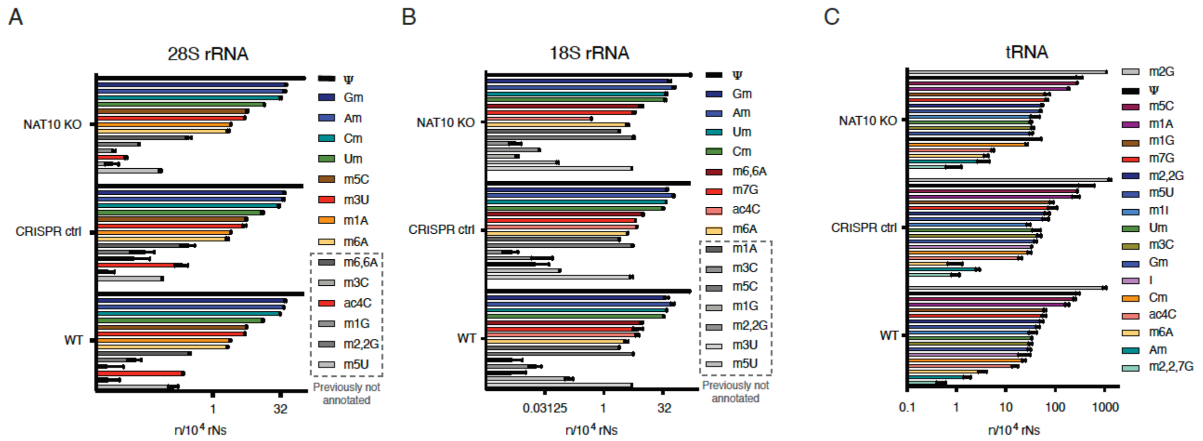


**Figure S2. RNA modification profiles validated by RNA MS in WT, NAT10 KO and CRISPR control cell lines.** a) 28S rRNA; b) 18S rRNA and c) tRNA. Previously not annotated RNA modifications are indicated with a dashed line. n=2 independent experiments.

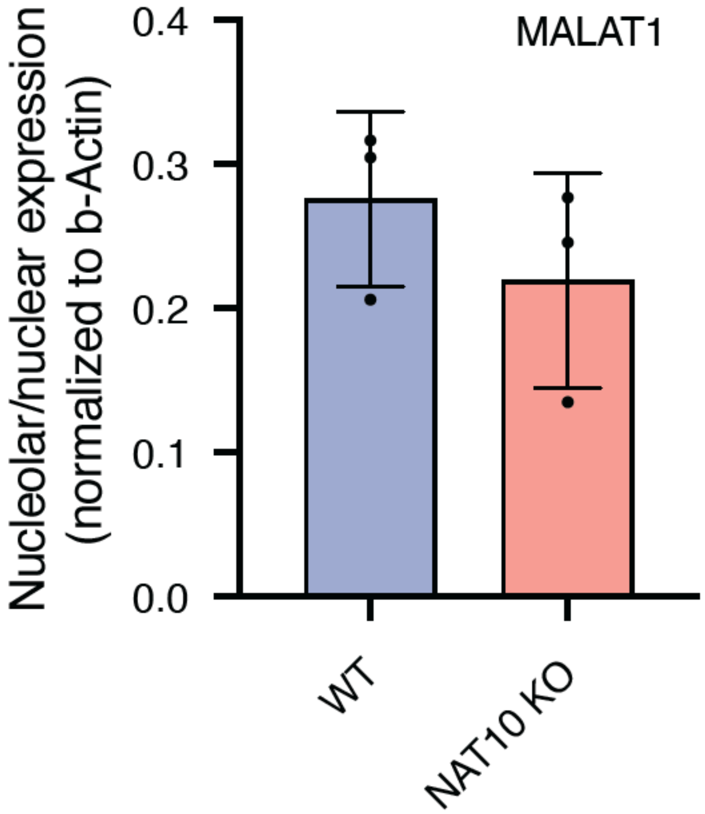


**Figure S3. MALAT1 localization in WT and NAT10 KO cells shown by qRT-PCR.** n=3 independent experiments. Data shown as average ± SD and individual data points are shown.
